# Improving Phenotypic Measurements in High-Content Imaging Screens

**DOI:** 10.1101/161422

**Authors:** D. Michael Ando, Cory Y. McLean, Marc Berndl

## Abstract

Image-based screening is a powerful technique to reveal how chemical, genetic, and environmental perturbations affect cellular state. Its potential is restricted by the current analysis algorithms that target a small number of cellular phenotypes and rely on expert-engineered image features. Newer algorithms that learn how to represent an image are limited by the small amount of labeled data for ground-truth, a common problem for scientific projects. We demonstrate a sensitive and robust method for distinguishing cellular phenotypes that requires no additional ground-truth data or training. It achieves state-of-the-art performance classifying drugs by similar molecular mechanism, using a Deep Metric Network that has been pre-trained on consumer images and a transformation that improves sensitivity to biological variation. However, our method is not limited to classification into predefined categories. It provides a continuous measure of the similarity between cellular phenotypes that can also detect subtle differences such as from increasing dose. The rich, biologically-meaningful image representation that our method provides can help therapy development by supporting high-throughput investigations, even exploratory ones, with more sophisticated and disease-relevant models.

## Introduction

Therapy development commonly ends in failure, potentially because it is too reliant on our incomplete understanding of the disease mechanism (Swinney and Anthony 2011; Sams-Dodd 2013). Phenotypic screening is an older approach to therapy development that is regaining momentum because of its hypothesis-free focus on correcting the disease and the sophisticated cellular models of disease now available. These new models of disease, including from a patient’s own cells, promise to better predict a candidate-therapy’s effect on the disease outcome in a patient if we can reliably measure their complex and subtle phenotypes. Improving our ability to predict the effect on disease outcome, even slightly, can dramatically increase the chance of successfully finding a therapy (Scannell and Bosley 2016).

High-Content Imaging (HCI) is a powerful technique to capture the diversity and subtlety of cellular phenotypes. A critical component of HCI, where the number of images may be in the hundreds of thousands, is developing an algorithm to automatically measure the cells. At present, a common HCI algorithm would target a simple phenotype such as cell death with as few as one or two expert-engineered image features (Singh et al. 2014b). This disconnect between the high information content of the image and the low information content of the typical analysis is why some still recommend manually reviewing all experimental images (Eggert 2013). In order to correctly prioritize a therapy candidate in an HCI screen by the biological relevance of its effect, we need to extract a more complete representation of the biology from each image.

Newly developed algorithms use deep neural networks to enable a data-driven approach to learning a representation of an image. The learned representation is the basis for the superior performance of deep learning over expert-engineered features across a variety of tasks (LeCun et al. 2015), but it commonly requires task-relevant labels (e.g. type of animal in the image for animal classification) on a large number of training examples. When many training examples are not available, an alternative is to use transfer learning, which starts with a neural network pre-trained on a different dataset and then trains the network on a dataset for the new task. The challenge of phenotypic screening is that we may not be able to induce all of the relevant phenotypes to gather training data, especially if we would like to capture a broad range of possible responses. Thus, transfer learning for HCI may be infeasible or impossible.

In this work we demonstrate accurate phenotypic discrimination without knowing the expected labels across a diverse set of phenotypes and within their subtle dose-response effects. We start with a Deep Metric Network that has learned to represent similar consumer images as nearby points in a continuous coordinate space, also referred to as an embedding. Without any additional training, we use a technique to dramatically improve our embedding for scientific images, that we have named Typical Variation Normalization, which enables us to surpass the present state-of-the-art for phenotype classification of an HCI screen and makes our results more robust to nuisance variation such as batch effect. We believe our work will enable a higher-level interaction between the scientist and their HCI dataset, capturing the range of complex and subtle cellular phenotypes of the disease model and placing them in a spatial relationship that is biologically meaningful.

## Results

For all of our studies, we use the BBBC021v1 dataset (Caie et al. 2010), available from the Broad Bioimage Benchmark Collection (Ljosa et al. 2012). The dataset represents the phenotype of the MCF-7 wild-type P53 breast cancer cell line after being exposed to a variety of chemical compounds. The phenotype is captured in a scientific image that is composed of three single-channel images that represent DNA, tubulin, and actin. To compare our method’s performance, we restrict our analysis to the compounds and concentrations described in Ljosa et al. (2013), where each compound at a concentration is referred to as a treatment. They define a subset of 103 treatments from 38 compounds that belong to one of 12 molecular mechanisms-of-action (MOA). As they note, only half of their MOA were identifiable visually, and the remainder were defined based on the literature.

## Overview of the approach

### Embedding Generation

We use a Deep Metric Network, based on the architecture described in Wang et al. (2014), that was pre-trained on a dataset of approximately 100 million RGB consumer images. The training task is to place images nearby in the embedding that are semantically similar, such as two images of the same type of lamp even if the camera angle, lamp color, and background environment are different. Each input image is shrunk or enlarged as necessary to fit the fixed input image size of ~200×200 pixels and produces a 64-dimensional embedding.

The input image into the network is a Cell Candidate Region, which is a cropped image that is centered on a cell (Fig 1A). We determine the centers of the cell candidates by segmenting only the DNA image to identify the positions of likely nuclei. We chose to segment the nuclei because they are relatively non-overlapping with excellent signal-to-background, making the segmentation simple and robust.

**Figure 1.**
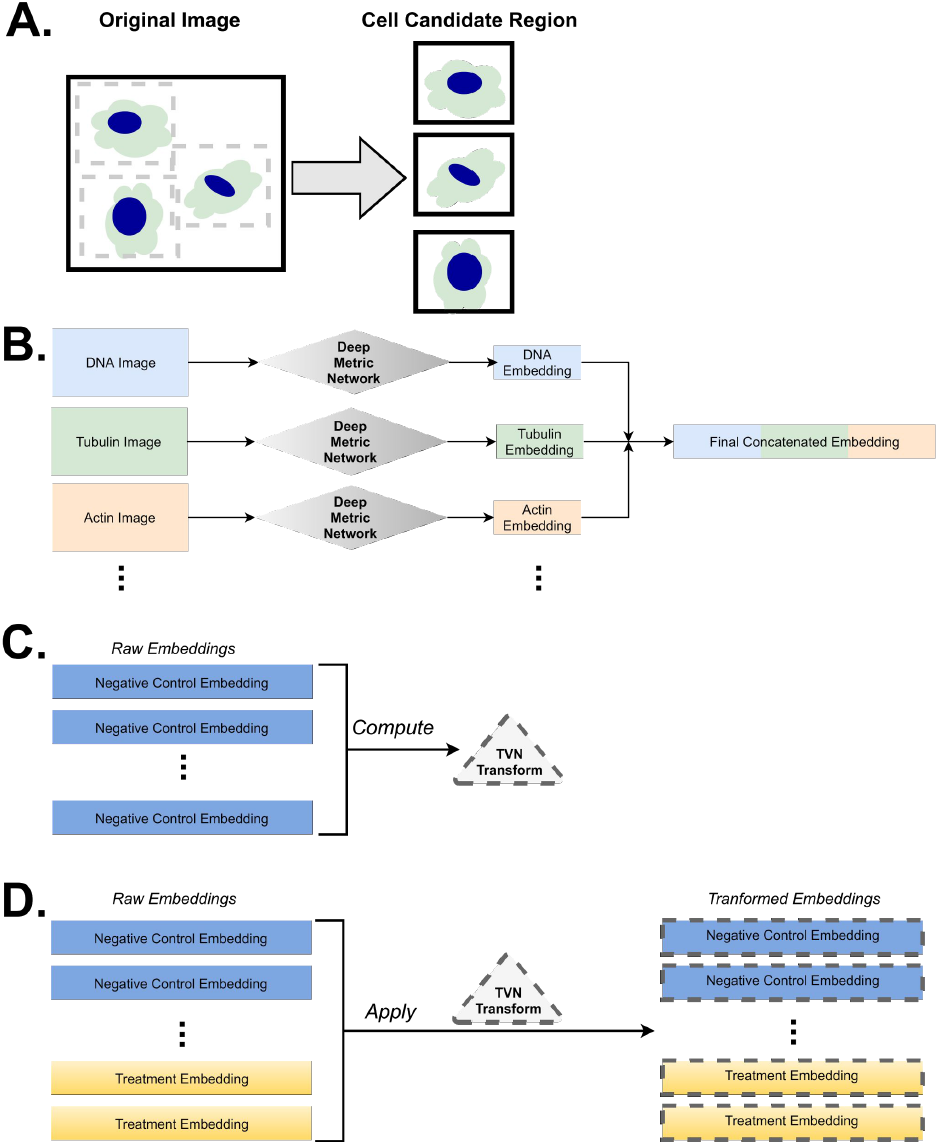
Overview of the approach. A scientific image is normally composed of several single-channel images. From the original image (1280 pixels ×1024 pixels), a patch (128 pixels ×128 pixels) is extracted that is centered on a Cell Candidate Region (A). To compute the embedding for a scientific image, we compute an embedding for each of the single-channel images using a pre-trained Deep Metric Network and then concatenate them (B). Typical Variation Normalization uses the collection of embeddings from the control conditions to compute a normalization transform (C) that is then applied to all of the control and experimental treatment embeddings (D).

The Deep Metric Network was not trained on scientific images, so we must convert them to match the bit-depth and RGB format of consumer images. Once converted, we feed each single-channel image into the Deep Metric Network to generate an embedding and then concatenate all of the single-channel embeddings into the final embedding (Fig 1B). This would result in a final embedding of 64 to 320-dimensions for a scientific image composed of 1 to 5 single-channel images, respectively. With the Cell Candidate Region approach, we have images from multiple cell locations (spread across wells and plates) for each treatment and thus have a collection of embeddings of each treatment. When comparing between treatments, we use the same Means method as Ljosa et al. (2013), which produces a single average value at each dimension, to generate a single embedding for each treatment.

### Typical Variation Normalization

The embeddings from the Deep Metric Network are not tuned to represent cellular phenotypes, but we can improve their representation by taking advantage of the common format of an HCI screen. Each plate in a screen has examples of the unperturbed cellular state in the negative control wells. The embeddings of the negative control sample the wide range of typical variation, arising from both methodology and biology. This typical variation provides two insights into how to improve the embeddings of cellular phenotypes: desensitize them to the axes of variation that separate unperturbed cells, and sensitize them to the axes of variation that subtly or rarely occur in unperturbed cells.

We capture the axes of variation using Principal Component Analysis (PCA) on the negative control embeddings without any dimensionality reduction. We then normalize each axis, which reduces the effect of axes with large variation and amplifies the effect of axes with little variation. We further reduce batch-to-batch variation by applying correlation alignment (Sun et al. 2016), aligning each batch with the entire experiment. All of these steps are calculated using only the negative control conditions, creating a transform for what we refer to as Typical Variation Normalization (Fig 1C). We then apply that same transform to all of the embeddings, both negative control and treated conditions (Fig 1D). This technique allows us to reshape the latent information of the embeddings from the Deep Metric Network to improve our representation of the cellular phenotypes.

### State-of-the-Art discrimination of biologically-similar phenotypes using a Deep Metric Network with Typical Variation Normalization

The current state-of-the-art method for classifying the BBBC021 phenotypes without additional training is the Factor Analysis method of Ljosa et al. (2013), which optimizes the ability of a small number of underlying factors to recreate the observed cell measurements. It achieves 94% accuracy for correctly identifying the MOA by matching with its nearest neighbor when using ~450 per-cell measurements with restrictions on matching the same compound (Not-Same-Compound or NSC). However, Pawlowski et al. (2016) note the Factor Analysis method has inconsistent performance between experiments, and the Means method, discussed above, is the preferred approach. Thus we also compare our results to the original Means method results of Ljosa et al. (2013) with 83% NSC accuracy, the best Means method with expert-engineered features of Singh et al. (2014a) with 90% NSC accuracy and the current state-of-the-art for the Means method of Pawlowski et al. (2016) with 91% NSC accuracy. Our Candidate Cell Region approach with Typical Variation Normalization (TVN) exceeds the current state-of-the-art performance with 96% NSC accuracy (Table 1). The accuracy for 9 of the 12 MOA is 100% with the majority of the error in classifying actin disruptors (Fig 2A). The ability of the Deep Metric Network embeddings with TVN to discriminate phenotypes is also evident from the two-dimensional visualization of the high-dimensional distance between embeddings as modeled by the t-Distributed Stochastic Neighbor Embedding (tSNE) technique (Maaten and Hinton 2008). MOA clusters are evident although there is not necessarily a unique cluster and outliers are present (Fig 2B). These results demonstrate the powerful generalizability of the Deep Metric Network with TVN, exceeding state-of-the-art at differentiating biological phenotypes *even though it was never trained on that data*.

**Table 1.** 1-Nearest Neighbor Mechanism-of-Action accuracy for BBBC021 treatments. TVN (Typical Variation Normalization).

|  |  | Not Same<br>Compound<br>Accuracy | Not Same<br>Compound or<br>Batch<br>Accuracy |
| --- | --- | --- | --- |
| **This work** | **TVN** | **96%**<br>**(99/103)** | **95%**<br>**(87/92)** |
| Pawlowksi<br>et al. (2016) | Means | 91%<br>(94/103) | Unable to<br>Calculate |
| Singh<br>et al. (2014a) | Means | 90%<br>(93/103) | 85%<br>(79/92) |
| Ljosa<br>et al. (2013) | Means | 83%<br>(86/103) | 58%<br>(53/92) |
|  | Factor<br>Analysis | 94%<br>(97/103) | 77%<br>(71/92) |

**Figure 2.**
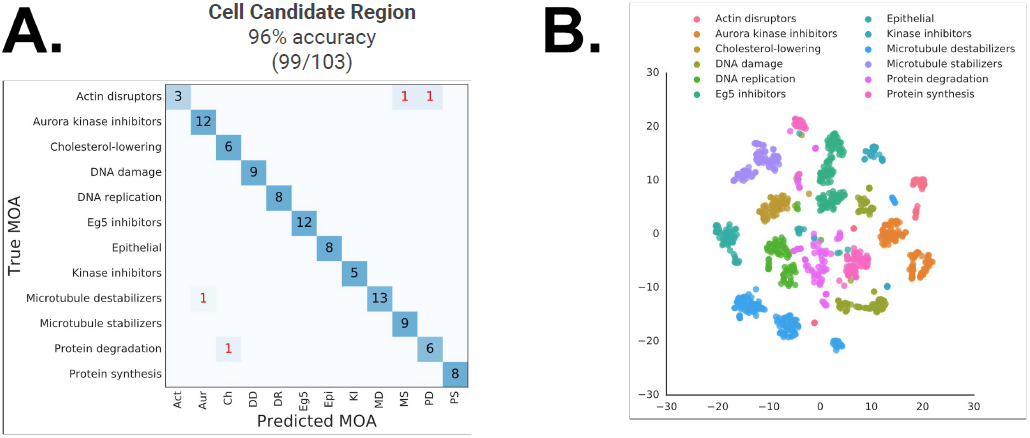
A Deep Metric Network with Typical Variation Normalization groups diverse phenotypes without any additional training. Confusion matrix for the 1-Nearest Neighbor prediction of treatments for Not-Same-Compound (A). Visualization of the embedding distance between each image using the t-Distributed Stochastic Neighbor Embedding (tSNE) algorithm to optimize the mapping from 192 dimensions to 2 dimensions for the the per-image average embedding (B).

### Nuisance variation may contribute to overestimation of BBBC021 phenotype classification accuracy

Within a screen, the layout of controls and experimental compounds is often more determined by constraints of automated liquid handling than by statistical concerns. As a result, nuisance variation, which we define as influence from batch, plate, or well-position effects, may give rise to undesired correlations between treatment measurements. These correlations can bias the results both towards or away from the correct classification when matching to ground-truth compounds. We see potential for bias from nuisance variation in the BBBC021 dataset from the small number of batches (1-3) that each MOA is spread across (S1 Fig) and the clustering of image embeddings from the same batch by tSNE (S2 Fig).

We can estimate the size of a method’s bias in a dataset by replacing the values for all treatments with the values from the nearest negative control well, in this case the negative control well that is located in the same row. Since we expect negative control conditions to be biologically equivalent, then we would expect the accuracy of the mock-treatment to be equal to random chance in the absence of bias. However, the NSC accuracy for the DMSO mock-treatments is higher than random chance for all of the approaches that we could evaluate (Table 2), reaching up to a three-fold higher odds for the Means of Ljosa et al. (2013). The prevalence of positive bias in the mock-treatments suggests that the NSC accuracy for the true treatments may also be overestimated due to nuisance variation.

**Table 2.**
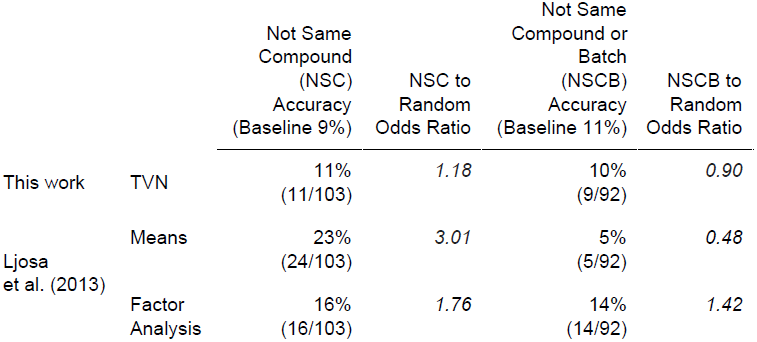
1-Nearest Neighbor Mechanism-of-Action accuracy for DMSO mock-treatments. TVN (Typical Variation Normalization).

### Deep Metric Network wsith Typical Variation Normalization maintains high phenotype classification accuracy after reducing the impact of nuisance variation

While completely removing the effect of nuisance variation is beyond the scope of this paper, we propose a simple extension to the 1-Nearest Neighbor benchmark that reduces the positive bias. In addition to preventing the predictions from matching to the same compound at a different concentration (NSC), we also restrict matching to any compound from the same batch (Not-Same-Compound-or-Batch or NSCB), reducing one of the strongest sources of measurement correlation. For NSCB analysis we must remove two MOAs (Cholesterol-lowering and Kinase-inhibitors) that are only present on a single batch, which leaves a total of 92 remaining treatments.

Comparing the NSCB and NSC performance on the DMSO mock-treatments, the NSCB accuracy is always closer to random chance than the NSC accuracy (Table 2). For the Means method of Ljosa et al. (2013) there may even be a flip to a negative bias. Thus reducing the batch effect with NSCB appears to create a less positively biased metric.

Assessing the NCSB performance on the true treatments, all approaches show a decrease in accuracy (Table 1), which further supports the idea that the NSC accuracy is an overestimate. The smallest drop in accuracy (96% to 95%) and highest overall performance is with the Candidate Cell Region approach with TVN. In contrast, the previous state-of-the-art Factor Analysis approach has a substantial drop in accuracy (94% to 77%), and even the more robust Means method of Singh et al. (2014a) has an NSCB accuracy difference of 10% with our approach. Thus, the advantage of our Deep Metric Network with TVN is even larger with a performance metric that reduces the positive bias from nuisance variation.

### Typical Variation Normalization improves separation of nuisance and phenotypic variation

The reduction in bias and improvement in classification performance with TVN is surprising since we do not perform dimensionality reduction to retain only the most informative dimensions. The PCA dimensions that capture a small amount of the total variation are often discarded as noise in other tasks, and increasing their impact through normalization could be interpreted as amplifying the noise. We demonstrate that, contrary to other tasks, the small total variation PCA dimensions are crucial for discriminating cellular phenotypes, while the large total variation PCA dimensions are actually more representative of nuisance variation.

We visualize the amount of information that each TVN dimension contains about the nuisance variation by studying the negative control embeddings. Since we compute the PCA transform on the negative controls, their embeddings have dimensions that are almost completely independent. This independence allows us to compute a per-dimension Mixed Effect model, where an embedding value for a Cell Candidate Region is the result of the fixed effects of batch and the individual effect of the image it was cropped from. Ordering the PCA dimensions by decreasing total variation, the largest magnitude of the batch effect is concentrated at the front of the embedding with the large total variation PCA dimensions (Fig 3A). This supports the idea that normalization of the large total variation PCA dimensions in the negative control helps to decrease the impact of nuisance variation.

**Figure 3.**
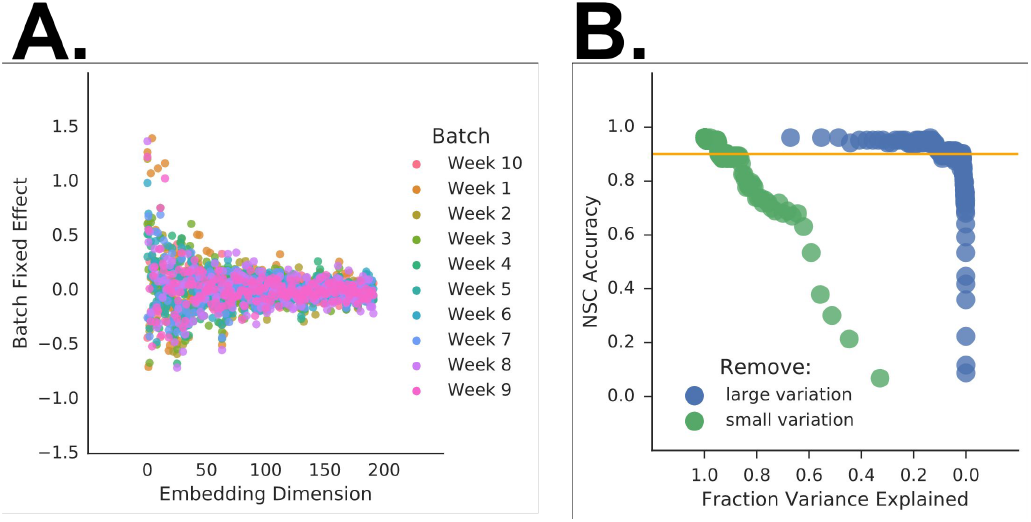
Typical Variation Normalization improves separation of nuisance and phenotypic variation. A Mixed Effect Model that calculates the contribution of nuisance variation to the embeddings of the negative controls shows that it is heavily weighted toward the front of the embeddingfor the batch effect (A). Performing dimensionality reduction by removing dimensions from either the “back” (small-variation removed first) or the “front” (large-variation removed first) demonstrates unequal effect on NSC accuracy (B). When removing from the back, the NSC accuracy quickly drops below 90% (orange line), even though the remaining dimensions capture still capture more than three quarters of the total variation. In contrast, removing from the front maintains an accuracy above 90%, even when the dimensions that remain capture less than a third of the total variation.

We evaluate the importance of the TVN dimensions for discriminating cellular phenotypes using ablation studies. We sequentially shorten the embedding by removing the dimension with the smallest total variation or the largest total variation. If we remove the final embedding dimensions with small variation first, we can have NSC accuracies well below 90% even though the embeddings capture more than 75% of the cumulative amount of variation (Fig 3B, green). Conversely, if we remove the the dimensions with large variation first, we can create embeddings that only capture ~20-30% of the cumulative amount of variation but still have NSC accuracies greater than 90% (Fig 3B, blue). These results indicate that the large variation PCA dimensions contain relatively little information, while the small variation PCA dimensions contain a lot of information that is relevant to discriminating cellular phenotypes. This is consistent with TVN desensitizing the embeddings to nuisance variation and sensitizing them to the relevant and subtle phenotypes of the cellular model.

### Deep Metric Embeddings can measure a dose-response across diverse phenotypes

Measuring the subtle changes in phenotype across a range of increasing dose can be critical for correctly prioritizing a therapy candidate. These measurements enable a more accurate determination of whether a candidate is active than a single high-dose, allow a candidate to be ranked by its potency, and provide insight into the structure/activity relationship for a candidate series (Inglese et al. 2006). We evaluated the ability of our approach to detect these subtle phenotypic changes within each of the diverse set of BBBC021 phenotypes.

If we compute phenotypic changes using the cosine distance as previously for MOA classification and the average DMSO embedding across all of the plates as the reference point, then all concentrations of each compound appear to be nearly equally distant (S3 Fig). We hypothesize that this is because the cosine distance has a bounded range, and the center of the DMSO conditions is far from any individual plate due to nuisance batch and plate effects. Since we cannot remove nuisance variation, we chose to increase the dynamic range of our measurements by using the logarithm of the euclidean distance, which does not have a bounded range and keeps the range of distances comparable across MOA.

With our approach we show that we can measure a dose-dependent phenotypic response across all of the MOAs (Fig 4). The average distance to DMSO negative control is far from zero (grey dashed line), supporting the idea that the center of the DMSO is far from any individual plate. However, for each of the twelve MOAs there is an increasing distance in each phenotype with increasing dose. For at least half of the MOA the dose-response curve also has the typical sigmoidal shape. This indicates that, even without training the network to recognize specific phenotypes, our Candidate Cell Region approach with TVN can capture the subtle phenotypic changes between small differences in dose.

**Figure 4.**
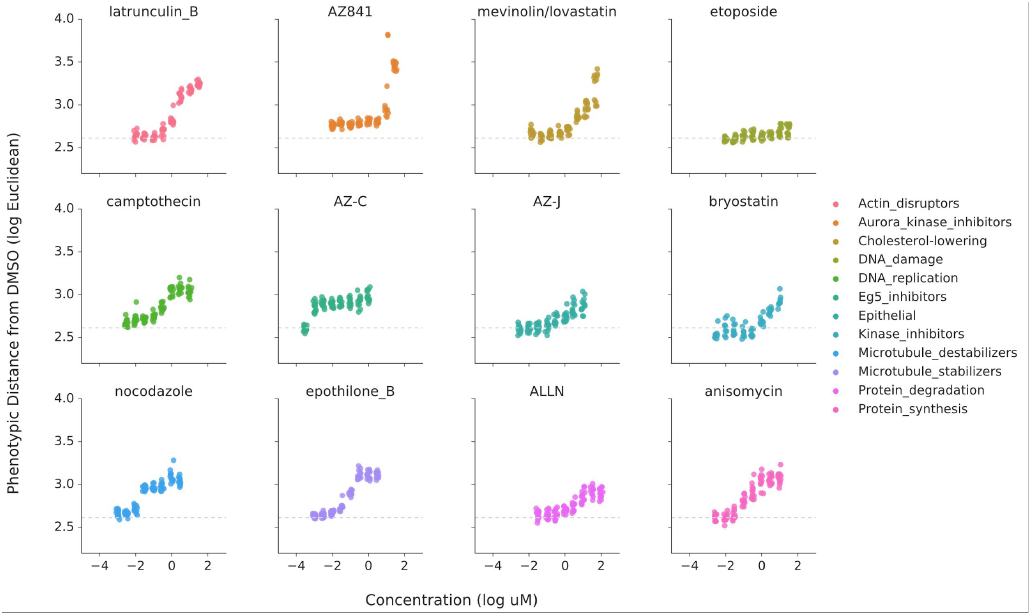
A Deep Metric Network with Typical Variation Normalization detects increasing phenotypic distance with increasing dose. We determined the phenotypic distance for each image by calculating the euclidean distance from each cell to the origin and then taking the average distance for the cells in an image. For the origin, we used the center of the negative control treatments. A representative example of a dose-response from each mechanism-of-action group is shown as well as the average distance from the negative controls to the center of the negative controls (grey dashed line).The shape of the curves range from a generally monotonic increase in distance with increasing dose (e.g. AZ841) to clearly sigmoidal (e.g. epothilone B and ALLN).

## Discussion

We demonstrate that a pre-trained Deep Metric Network with Typical Variation Normalization can accurately, robustly, and sensitively differentiate across a diverse set of phenotypes without any additional training. Our approach can beat the present state-of-the-art phenotypic classification, even when using a more challenging metric for accuracy. The power of our method is that we improve our representation of the biology by taking advantage of the tremendous variation in the negative controls, which is normally a source of experimental and statistical frustration.

The ability of control cells to provide insight into cellular variation was also noted in Ljosa et al. (2013). Their Factor Analysis method on expert-engineered features performed equally well when calculated on control or treated cells. However, their interpretation, and thus implementation, differs significantly from ours. They focused on finding the few fundamental modes to explain their measures and considered variations in single measures as noise. This is consistent with their dimensionality reduction from ~450 features to 50 latent factors for their peak performance. They also perform normalization as an initial step on their raw measures in order to reduce plate-to-plate nuisance variation rather than on the latent factors. In contrast, we take advantage of the concise semantic embedding from the Deep Metric Network. We interpret every variation that occurs within the negative control cells as potentially meaningful, and we perform normalization on our transformed dimensions in order to both reduce the impact of nuisance variation and increase the impact of subtle or rare phenotypes.

The power of deep learning with supervised training is shown in a recent paper by Kraus et al (2016). Training on 15% of the images from the BBBC021 ground-truth labels, they demonstrated the ability to simultaneously learn how to classify the images and segment the cells. Interestingly, their highest accuracy of 97% on the BBBC021 dataset is only slightly better than the 96% we achieve without any training on the data. Similarly, Godinez et al. <u>(2017)</u> used supervised training on BBBC021 with thirteen classes (twelve MOA and DMSO) to classify the MOA across a dose-response curve, demonstrating the ability reasonably estimate concentrations of half-maximal response. We did not pursue a similar supervised approach for three reasons. The first is that we wanted our evaluation to be similar to an actual screen where it is not feasible to train on all of the possible phenotypes. The second is that training would require splitting the BBBC021 dataset into training and testing subsets, making it impossible to evaluate our performance identically to Ljosa et al (2013). The third is that the small size of the BBBC021 dataset makes it very easy to overfit, leading to an overestimation of the classification accuracy. However, if one can gather training data for a task that covers the complete set of potentially relevant phenotypes, then supervised deep learning may be the superior approach.

While we believe our results are promising, we recognize that there are several limitations that must be taken into account. Our analysis is presently limited to a relatively small screen of a single cancer cell line where nuisance factors are likely biasing the results. In future work, we will investigate how the performance of our approach generalizes across a variety of cellular models and fluorescent markers and on larger datasets with more ground-truth examples, ideally with more randomization in the layout of conditions. We also cannot compare our NCSB metric across all of the previously published results, particularly Pawlowksi et al. (2016) that used a different neural network architecture. Our future work will investigate how other deep learning architectures perform using the Cell Candidate Region with TVN. Lastly, we do not have ground-truth for the correct shape of the dose-response curve for each compound. In future work, we will further investigate the validity of the dose-response shape in our TVN embedding and, in particular, the sensitivity of our approach to detecting compound effects at low doses.

Our approach places the images of an HCI screen within an embedding where distance between points meaningfully captures differences between cellular phenotypes. This allows us to condense the results of many thousands of images into a single plot. Scientists will no longer need to pick a simple phenotype, such as cell death, because it is easy to quantify. Our goal is to enable a scientist to visualize and interact with their image dataset with our embedding as a starting point for finding patterns, developing hypotheses, and investigating interesting outliers. We hope that this will increase the adoption of more sophisticated and disease-relevant phenotypic models for screens (Vincent et al. 2015; Caicedo et al. 2016), leading us to a more successful therapeutic pipeline.

## Materials and Methods

### BBBC021 Image Dataset

We use the BBBC021v1 dataset (Caie et al. 2010), available from the Broad Bioimage Benchmark Collection (Ljosa et al. 2012). The dataset uses the MCF-7 wild-type P53 breast cancer cell line, and each image is composed of three single-channel images: DAPI/DNA, actin, and tubulin. A total of 55 96-well plates were imaged in batches of 3-6 plates over 10 weeks. Each plate has 6 negative control (DMSO) wells and 6 positive control (Taxol) wells. Each compound is tested at 8 concentrations on triplicate plates where each compound and concentration pair is referred to as a treatment. The entire dataset has 113 compounds for a total of 906 treatments. The Ljosa et al. (2013) subset is 38 compounds for a total of 103 treatments.

### Embeddings from the Deep Metric Network

#### Image Preprocessing

We convert all of the 16-bit integer images into 32-bit float images, so we can perform our preprocessing without introducing errors from integer rounding. We compute a flatfield image for each plate and channel similar to Singh et al. (2014a) except we use the 10th percentile rather than the median, and we blur the resulting image with a gaussian sigma of 50.

Next, we divide each image by its appropriate flatfield image, producing an image that represents the signal/background at each pixel location. To improve the dynamic range before converting to an 8-bit integer image, we set the minimum value to 1.0 and take the natural log of each pixel value. We then find the minimum and maximum values of the image, and we clip the maximum to be no greater than 5 to prevent extreme values. We then linearly re-scale the minimum and maximum to be the values 0 and 255. To create the final 8-bit RGB image, we round the floating point values, convert the values to 8-bit unsigned integer, and place the same image into each of the R, G, and B channels.

#### Network Architecture

The Deep Metric Network used is built and trained similar to Wang et al. (2014) with a triplet-loss function for both a visual and semantic task although we only use the semantic embedding. Briefly, the training for the semantic task used a dataset generated from click-data from text queries. Images that were often clicked on for the same query were considered semantically similar. The input images were of variable size, so each input image is scaled to a final dimension of 224 pixels × 224 pixels.

#### Cell Candidate Regions

Cell Candidate regions are a 128×128 pixel patch with the patch centered on a nucleus. Cells with nuclei centers closer than 64 pixels from the edge are excluded so that the image patch is always fully within the boundary of the image.

For all comparisons to previous results, we start with the DAPI center locations of Ljosa et al. (2013), and filter out any that are too close to the border. For the dose-response evaluation, no nuclei centers were defined for concentrations that were not used in Ljosa et al. (2013), so we computed our own nuclei centers for all wells. Briefly, the signal/background processed image described in the Image Processing section is binarized into foreground and background regions using Otsu’s method (Otsu 1979). Two rounds of binary erosion and propagation are applied to the resulting mask and all connected components in the mask are identified. After filtering components with area <200 pixels, a watershed segmentation is applied to each component to identify candidate cells. Initial candidate cell center are placed at the center of mass of each resulting cell label. To ameliorate over-segmentation, candidate cell centers are pruned so that no two centers are closer than 20 pixels to each other.

### Typical Variation Normalization

With all the negative controls in the experiment, we compute the PCA basis and the per-dimension normalization to zero-center and unit variance. We then apply this transform and whitening to the embeddings of both the control and experimental treatment. Next, we perform correlation alignment as in Sun et al. (2016). On the transformed negative controls, we compute the covariance matrix of the entire experiment and a covariance matrix per-batch. We then align both the control and experimental treatments in each batch, using the covariance matrix of all negative controls as the target and the covariance matrix of each batch of negative controls as the source.

### 1-Nearest Neighbor Mechanism-of-Action Assignment

We perform 1-Nearest Neighbor classification as in Ljosa et al. (2013). First, all of the embeddings for a treatment on a plate are averaged and then the median across the three plates is used to create a single embedding for the treatment. To determine the distance between embeddings, we use the cosine distance. To compare with previous results, we eliminate any match to the same compound (Not-Same-Compound or NSC). We also extend the analysis to include not matching anything on the same batch (Not-Same-Compound-or-Batch or NSCB).

### Visualization of Embeddings

To create a single embedding for each image, we take the mean of all cell embeddings within the image. We then compute the pairwise distance matrix of images using the cosine metric. This distance matrix is then transformed into a two dimensional representation with tSNE on the default settings of Scikit-Learn (Pedregosa et al. 2011).

### DMSO Mock-Treatments

To create mock-treatments, we select the DMSO well that is in the same row as the treatment and take the mean of all of the cell measurements. Since we did not have access to the well-level measurements from Singh et al. (2014a) and Pawlowksi et al. (2016), the were excluded from this analysis, and we used the Means approach of Ljosa et al. (2013) instead. All concentrations of the compound are in the same row and thus receive the same mock-treatment, but they will not be matched by either NSC or NSCB. The 1-NN classification is then computed as above.

The odds for random chance are calculated as follows. The number of correct by random chance is calculated using the fraction correct based on random matching of MOA multiplied by the total number of treatments. The number of incorrect by random chance is the total number of treatments minus the number of correct by random chance, and the odds are correct/incorrect.

### Mixed Effect Model of Batch Variation

To compute the contribution of batch to the embeddings, we restrict our analysis to the negative controls, and we calculate a mixed effect model using the “lmer” function in the “arm” package (Gelman and Hill 2007). The model was fit with batch as a fixed effect and a unique identifier for the image as the random effect.

### Dose-Response Distance

Although we compute our own nuclei centers as above, we restrict the analysis to compounds that were defined in Ljosa et al. (2013) but extend to all concentrations rather than just the concentrations defined as active. We perform Typical Variation Normalization and compute the distance to each cell using the euclidean distance to the center of the DMSO negative controls. The distance for each image is then the mean of all of its cell distances.

## Acknowledgements

*We would like to thank members of the Google Accelerated Science team for their thoughtful discussion and feedback on early drafts of this paper, in particular Geoff Davis, Michelle Dimon, Steven Kearnes, Jason Miller, Philip Nelson, Annalisa Pawlosky, Patrick Riley, and Samuel Yang.*

*We would also like to thank Anne Carpenter, Neil Carragher, Nick Pawlowski, and Shantanu Singh for their generous aid in understanding the BBBC021 dataset and helpful discussion of their analyses.*

---

**Supplemental Figure 1.**
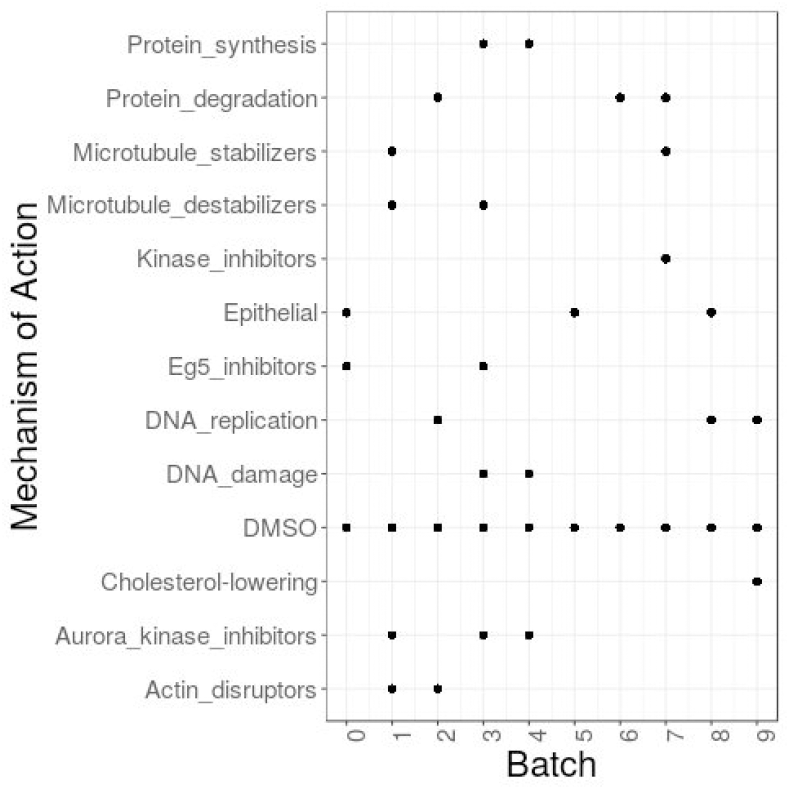
Compounds with the same mechanism-of-action are not evenly spread across imaging batches of BBBC021.

**Supplemental Figure 2.**
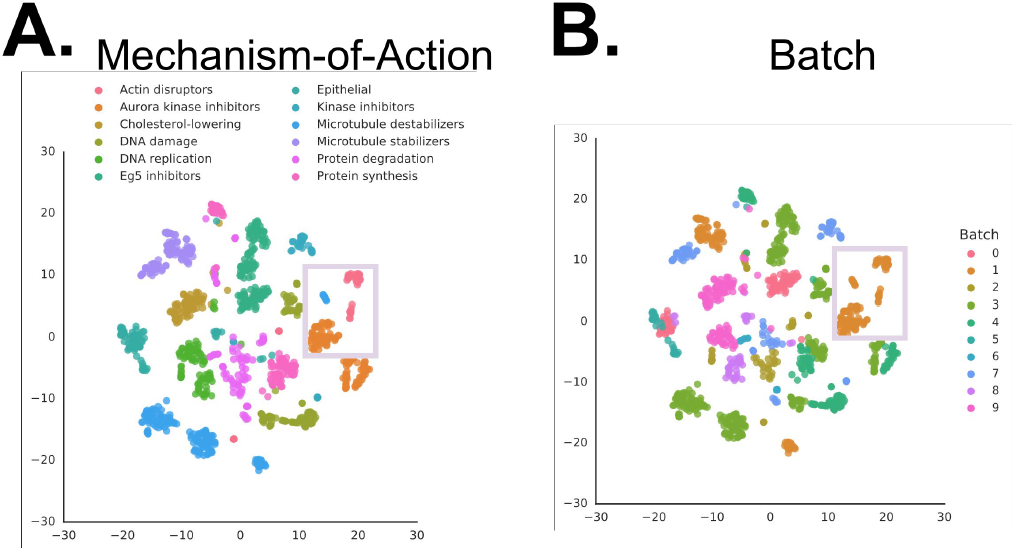
Candidate Cell Region embeddings of BBBC021 cluster near treatments from the same batch. Visualization of the embedding distance between each image using the t-Distributed Stochastic Neighbor Embedding (tSNE) algorithm to optimize the mapping from 192 dimensions to 2 dimensions for the the per-image average embedding with color labels for mechanism-of-action (A) or batch (B). Placement of nearby images in the embedding reflect mechanism-of-action as well as batch (light grey square).

**Supplemental Figure 3.**
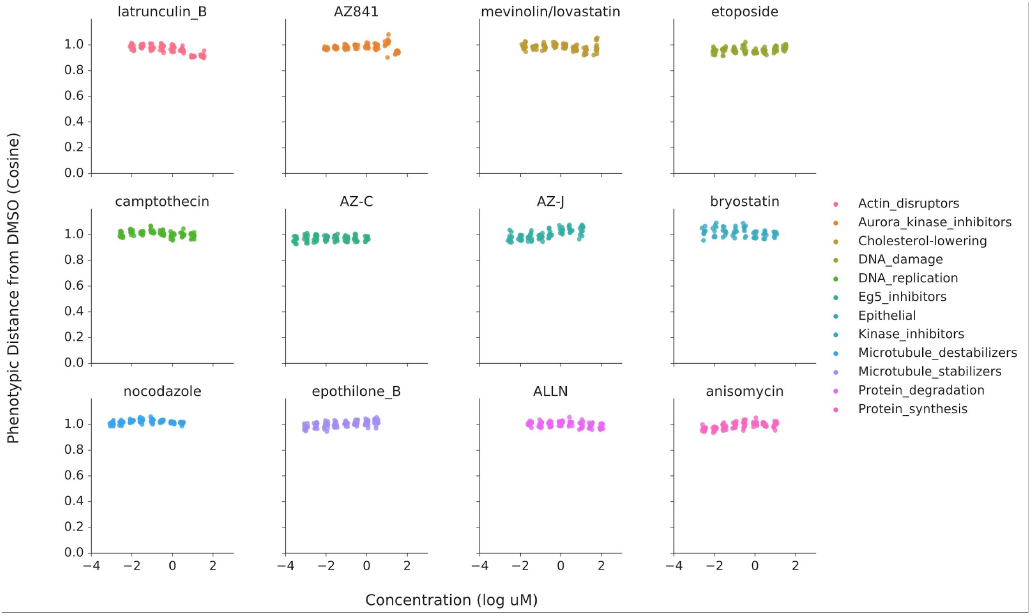
Cosine distances from the DMSO center are all nearly equally distant.

